# Phosphorylation of Arp2 is not essential for Arp2/3 complex activity in fission yeast

**DOI:** 10.1101/372706

**Authors:** Alexander E. Epstein, Sofia Espinoza-Sanchez, Thomas D. Pollard

**Affiliations:** Departments of Molecular Cellular and Developmental Biology, Yale University, PO Box 208103, New Haven, CT 06520-8103 USA; Departments of Molecular Biophysics and Biochemistry, Yale University, PO Box 208103, New Haven, CT 06520-8103 USA; Department of Cell Biology, Yale University, PO Box 208103, New Haven, CT 06520-8103 USA

## Abstract

LeClaire *et al.* presented evidence that phosphorylation of three sites on the Arp2 subunit activates Arp2/3 complex to nucleate actin filaments. We mutated the homologous residues of Arp2 (Y198, T233 and T234) in the fission yeast genome to amino acids that preclude or mimic phosphorylation. Arp2/3 complex is essential for the viability of fission yeast, yet strains unable to phosphorylate these sites grew normally. Y198F/T233A/T234A Arp2 was only nonfunctional if GFP-tagged, as observed by LeClaire *et al.* in *Drosophila* cells. Replacing both T233 and T234 with aspartic acid was lethal, suggesting that phosphorylation might be inhibitory. Nevertheless, blocking phosphorylation at these sites had the same effect as mimicking it: slowing assembly of endocytic actin patches. Mass spectrometry revealed phosphorylation at a fourth conserved Arp2 residue, Y218, but both blocking and mimicking phosphorylation of Y218 only slowed actin patch assembly slightly. Therefore, phosphorylation of Y198, T233, T234 and Y218 is not required for the activity of fission yeast Arp2/3 complex.

**Summary:** Previous research concluded that phosphorylation at three sites on Arp2 is necessary to activate Arp2/3 complex. Epstein et al. make genomic substitutions blocking or mimicking phosphorylation to demonstrate that phosphorylation of these three sites does not regulate Arp2/3 complex in fission yeast.

## INTRODUCTION

Assembly of branched actin filament networks drives cellular processes including cell motility and clathrin-mediated endocytosis (Blanchoin et al., 2014; Weinberg and Drubin, 2012). The seven-subunit Arp2/3 complex builds these networks by binding to the side of a “mother” actin filament and nucleating a “daughter” filament branch (Mullins et al., 1998). Activation of Arp2/3 complex depends on binding of nucleation-promoting factors (NPFs) (Machesky and Insall, 1998; Machesky et al., 1999; Rohatgi et al., 1999; Winter et al., 1999; Yarar et al., 1999) which induce a conformational change (Espinoza Sanchez et al., 2017, preprint; Hetrick et al., 2013) and promote binding of the complex to the side of a mother filament (Ti et al., 2011). For example, the NPF Wiskott-Aldrich syndrome protein (WASp) is recruited to sites of endocytosis where it activates Arp2/3 complex (Winter et al., 1999). Arp2/3 complex then builds a “patch” of branched actin filaments that generates force to internalize endocytic vesicles (Carlsson and Bayly, 2014). In motile cells, the SCAR/WAVE complex activates Arp2/3 complex along the leading edge of the cell, stimulating the formation of the lamellipodium that sweeps the cell forward (Insall and Machesky, 2009). Regulation of Arp2/3 complex is essential to control the localization and assembly of branched actin networks.

LeClaire *et al.* discovered that purified *Acanthamoeba* Arp2/3 complex lost its ability to nucleate actin filaments when treated with serine/threonine and tyrosine phosphatases (LeClaire et al., 2008). Furthermore, antibodies to phosphothreonine and phosphotyrosine interacted with the Arp2 and Arp3 subunits of Arp2/3 complex from *Acanthamoeba*, *Bos taurus* and humans. Mass spectrometry was used to identify phosphorylation of the highly conserved T237 and T238 residues of amoeba Arp2. The location of Y202 near these threonines suggested that it might also be phosphorylated. LeClaire *et al.* investigated the role of phosphorylation at Y202, T237 and T238 in regulating the *Drosophila* Arp2/3 complex. Depletion of Arp2 compromised the formation of lamellipodia in *Drosophila* S2 cells. This defect was rescued by expression of wild-type Arp2-GFP, T237A/T238A Arp2-GFP or Y202A Arp2-GFP, but not by Y202A/T237A/T238A Arp2-GFP, indicating that phosphorylation of either the two threonines or the tyrosine is essential for Arp2/3 complex activity. A *Drosophila* kinase that phosphorylates these residues has been identified: in 2015, LeClaire *et al.* reported that the Nck-interacting kinase (NIK) can phosphorylate several Arp2/3 complex subunits, including Arp2 at Y202, T237 or T238 (LeClaire et al., 2015). NIK restored the actin nucleation activity of purified Arp2/3 complex after the complex was inactivated by treatment with serine/threonine and tyrosine phosphatases.

LeClaire *et al.* originally suggested that phosphorylation at Y202, T237 and T238 activates the Arp2/3 complex by disrupting inhibitory interactions of these residues with R409 of the Arp3 subunit and R105 and/or R106 of the ARPC4 subunit (LeClaire et al., 2008). A 2011 study employed molecular dynamics (MD) simulations to study the effects of the interactions involving these phosphorylated residues on the structure of the Arp2/3 complex (Narayanan et al., 2011). During all-atom MD simulations of native Arp2/3 complex for 30 ns, Arp2 shifted 3-4 Å relative to Arp3 from its position in the inactive crystal structure (Robinson et al., 2001) towards the short-pitch actin helix in the branch junction (Rouiller et al., 2008). This shift was about 2-fold larger when either T237 or T238 of Arp2 was phosphorylated and/or R105 of ARPC4 was replaced with alanine, although the changes during the simulation time explored were far short of the 30 Å displacement of these subunits in the branch junction. As predicted by the MD simulation results, substituting alanine for R105 and R106 partially activated purified Arp2/3 complex without an NPF (Narayanan et al., 2011).

These papers make a strong case that Arp2 phosphorylation relieves autoinhibitory interactions between two threonines or a tyrosine of Arp2 and three arginines on the Arp3 and ARPC4 subunits, inducing a conformational change that partially activates the complex. However, other evidence indicates that phosphorylation at the three proposed sites is not essential for some activities of Arp2/3 complex. For example, replacing the two threonines and the tyrosine with alanine in in *Dictyostelium* Arp2 slowed development but not pseudopod extension in a chemotaxis assay (Choi et al., 2013). Purified *Bos taurus* Arp2/3 complex can nucleate one actin branch per complex under ideal conditions *in vitro* (Higgs et al., 1999), but electron density maps of a 2.0 Å crystal structure of bovine Arp2/3 complex from the same preparation (Robinson et al., 2001) had no density corresponding to phosphorylation of Y202, T237 or T238 (Fig. S1 A). Therefore, further study was needed to determine if phosphorylation at the proposed sites regulates the Arp2/3 complex.

Here we explore the effect of Arp2 phosphorylation on Arp2/3 complex activity in the fission yeast *Schizosaccharomyces pombe,* where efficient homologous recombination facilitates making mutations in the genome (Bahler et al., 1998; Grimm et al., 1988) and quantitative fluorescence microscopy assays are available to characterize actin assembly during endocytosis (Berro and Pollard, 2014a). The proposed phosphorylation sites are conserved in *S. pombe* as Y198, T233 and T234 (Fig. S1 B). We made Arp2 mutations in the fission yeast genome that block phosphorylation or mimic constitutive phosphorylation at these three sites. Arp2 is essential for viability in *S. pombe* (Morrell et al., 1999) and therefore any mutations that inactivate Arp2/3 complex should be lethal. We determined whether strains with each Arp2 mutation were viable and measured the rate of actin patch formation in each viable strain to investigate how phosphorylation at each site would affect Arp2/3 complex activity.

## RESULTS AND DISCUSSION

### Detecting Arp2 phosphorylation with mass spectrometry

Mass spectrometry of Arp2/3 complex purified in the presence of phosphatase inhibitors revealed phosphorylation of Arp2 at Y198 and Y218, a previously unexplored but widely conserved site in eukaryotes (Fig. S1, C and D). We did not detect phosphorylation of T233 or T234. This does not rule out their phosphorylation *in vivo*, since phosphorylation of Arp2 at Y198 and Y218 was lost in *S. pombe* Arp2/3 complex purified without phosphatase inhibitors. It is possible that the addition of phosphatase inhibitors was insufficient to prevent dephosphorylation at T233 and T234.

### Generation and viability of Arp2 mutant fission yeast strains

To determine the effect of phosphorylation at the sites identified by LeClaire *et al.* on Arp2/3 complex activity in fission yeast, we generated haploid *S. pombe* strains with all possible combinations of mutations that preclude or mimic phosphorylation at Y198, T233 and T234 (Fig. 1 A; and Fig. S1 G). Alanine was used to preclude threonine phosphorylation, and aspartic acid to mimic it. We used phenylalanine and glutamate to prevent and mimic phosphorylation at Y198, respectively, because of their greater structural similarity to tyrosine.

**Figure 1.**
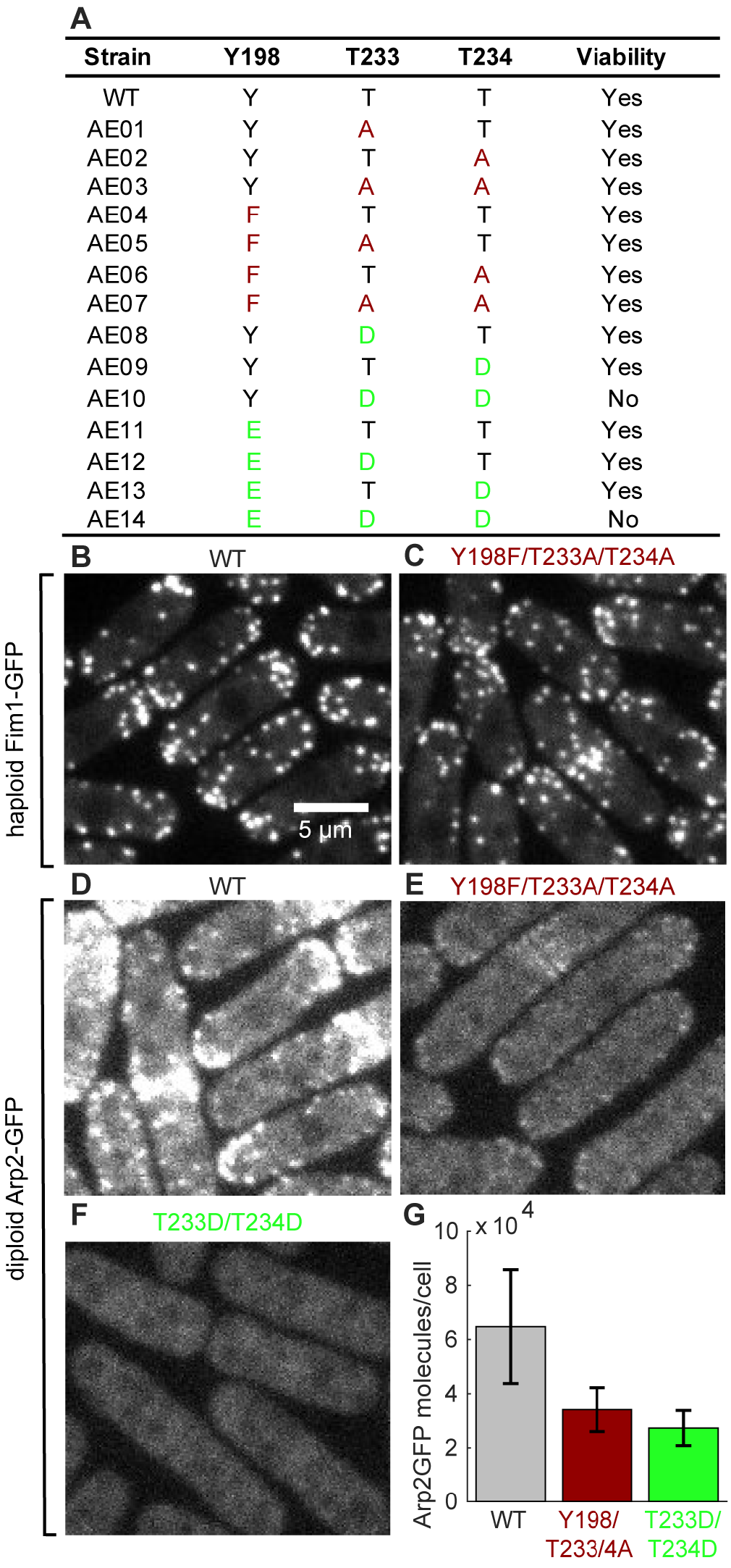
*S. pombe* strains with mutations of Arp2 residues Y198, T233 and T234. **(A)** Viability of haploid strains with mutations blocking or mimicking phosphorylation of residues Y198, T233 or T234 of Arp2, measured by growth of tetrads on YE5S plates at 25°C. Dark red: mutations blocking phosphorylation; light green: mutations mimicking it. **(B-F)** Sum projection confocal fluorescence images (6 z-sections with 0.6 μm spacing). **(B and C)** Haploid *S. pombe* expressing Fim1-GFP with **(B)** wild-type Arp2 or **(C)** Y198F/T233A/T234A mutant Arp2. (D-F) Diploid *S. pombe* strains with one copy of untagged wild-type Arp2 and one copy of **(D)** wild-type Arp2-GFP, **(E)** Y198F/T233A/T234A Arp2-GFP or **(F)** lethal mutant T233D/T234D Arp2-GFP. **(G)** Measurements of Arp2-GFP molecules per cell in diploid strains (mean ± SD, n = 47-82)

All strains with mutations preventing phosphorylation were viable, including the triple Arp2 Y198F/T233A/T234A mutant (Fig. 1A), even though Arp2/3 complex activity is necessary for viability of fission yeast (Morrell et al., 1999). Furthermore, cells depending on the triple mutant Arp2 successfully assembled actin patches marked by the actin crosslinking protein Fim1-GFP (Fig. 1 C; and Fig. 2 C). Therefore, Arp2 phosphorylation at Y198, T233 and T234 is not essential for Arp2/3 complex activity in *S. pombe*.

**Figure 2.**
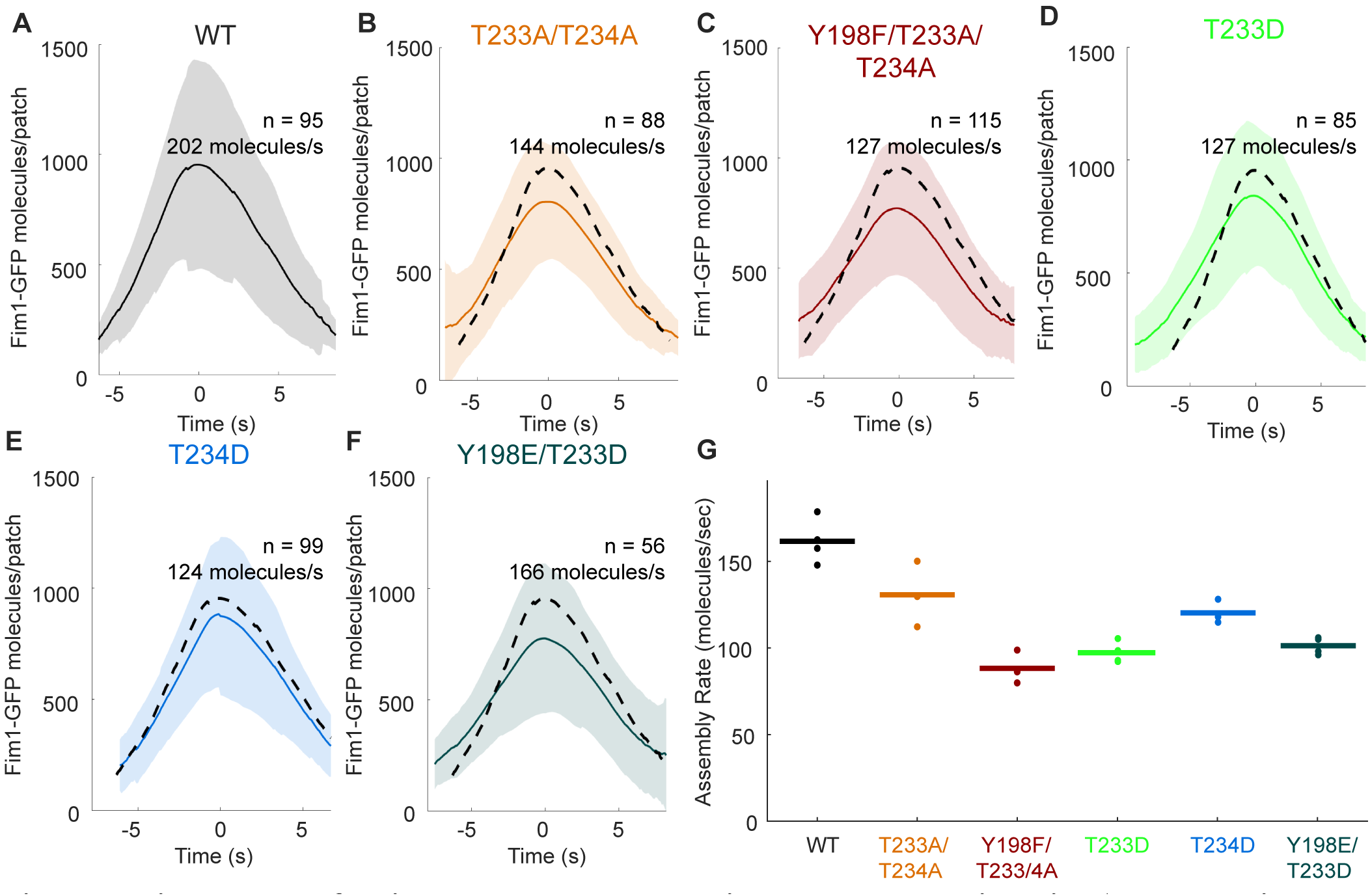
Time course of actin patch assembly and disassembly by strains with Arp2 mutations either blocking or mimicking phosphorylation at Y198, T233 and T234. **(A-F)** Mean numbers of Fim1-GFP molecules over time in 56-115 aligned actin patch tracks from haploid *S. pombe* strains with **(A)** wild-type Arp2, **(B and C)** Arp2 with mutations blocking phosphorylation, and **(D-F)** Arp2 with phosphomimetic mutations. Shaded regions indicate standard deviations. Dashed lines represent mean numbers of molecules per patch from the wild-type strain in Panel A. **(G)** Rates of patch assembly for 3-4 replicates (27-115 patches per replicate) of time lapse movies of *S. pombe* with wild-type Arp2 and Arp2 with mutations blocking or mimicking phosphorylation. Points indicate assembly rates of each replicate; horizontal bars denote the mean assembly rates.

By contrast, when a C-terminal GFP tag was added to the Arp2 subunit with the Y198F/T233A/T234A triple mutation, the strain was not viable. This parallels the failure of Y202A/T237A/T234A Arp2-GFP to rescue lamellipodial assembly in *Drosophila* cells depleted of Arp2 (LeClaire et al., 2008).

Most strains with mutations mimicking phosphorylation of Y202, T233, and T234 in Arp2 were also viable, although the two strains with the T233D mutation displayed a growth defect at high temperatures (Fig. S1 E). However, the two strains with the double T233D/T234D mutation to mimic phosphorylation at both threonines (AE10 and AE14) were not viable (Fig. 1 A). We assume that the T233D and T234D substitutions mimic phosphothreonine as aspartic and glutamic acid mimic phosphoserine, phosphthreonine and phosphotyrosine in other proteins (Dephoure et al., 2013). For example, the biochemical properties of cofilin with the S3D mutation are similar to those of phosphorylated cofilin (Blanchoin et al., 2000). Phosphomimetic substitutions are imperfect and can fail, especially at tyrosine residues because of the structural dissimilarity between phosphotyrosine and glutamate (Anthis et al., 2009). However, failure is most likely if the phosphorylated residue must fit precisely into a binding site, for example on an adapter protein (Dephoure et al., 2013). This is not likely to pose an issue at T233 and T234, as they are located on the interior of the Arp2/3 complex (Robinson et al., 2001), and phosphorylation at these sites is proposed to disrupt binding interactions between Arp2 and ARPC4 (LeClaire et al., 2008).

### Effect of Arp2 mutations at proposed phosphorylation sites on actin patch assembly

We used quantitative fluorescence microscopy of live cells (Berro and Pollard, 2014a) to compare the rates at which wild-type and mutant Arp2/3 complex assembled branched actin filaments in endocytic patches labeled with Fim1-GFP. Since individual actin patches (Fig. S2 A) vary in their total fluorescence and timing of events (Fig. S2 B), we used continuous alignment to average the assembly and disassembly curves of 27-115 patches (Fig. S2 C). We calculated the rate of actin patch assembly from the slope of the assembly phase of the averaged and aligned data.

All of the viable haploid strains depending on Arp2/3 complex with mutations that prevent or mimic phosphorylation of Arp2 residues Y198, T233 or T234 assembled and disassembled actin patches (Fig. 2), but all mutations of the three residues decreased the rate at which Fim1-GFP accumulated in the patches (Fig. 2 G; Fig. S2 G). Cells depending on Arp2 with the triple Y198F/T233A/T234A mutation assembled actin patches more slowly than cells with T233A/T234A Arp2. Actin patches assembled slower in the strains with the Arp2-T233D phosphomimetic mutation, which displayed a growth defect at 36°C, than in those with the Arp2-T234D mutation, which grew normally.

The subtle phenotypic changes that we observed in fission yeast strains with mutations of Arp2 phosphorylation sites are consistent with those detected in previous studies on other cells. On one hand, mutating Arp2 phosphorylation sites can produce strong phenotypes: *Dictyostelium* cells with the Arp2 Y202F/T237A/T238A mutations develop very slowly when faced with starvation (Choi et al., 2013). On the other hand, these mutant cells had only subtle defects in speed and directionality during chemotaxis, a short-lived process much more akin to endocytosis. Y202A/T238A/T238A Arp2-GFP did not rescue lamellipodial assembly in cultured *Drosophila* cells depleted of Arp2 (LeClaire et al., 2008), but we found that the severity of this phenotype is likely due to the presence of the C-terminal GFP tag on the mutated Arp2.

If phosphorylation of Y198, T233 and T234 plays a role in regulating the Arp2/3 complex, then we would expect blocking and mimicking phosphorylation at these sites to have opposite effects on Arp2/3 complex activity. However, mutations that prevent phosphorylation and mutations that mimic it both modestly decreased the rate of endocytic actin patch assembly in yeast cells. Therefore, the Arp2 mutations we made compromised assembly by some other mechanism, as explored in the next section.

### Observations of mutated Arp2-GFP in diploid cells

To determine how lethal mutations such as T233D/T234D Arp2 and Y198F/T233A/T324A Arp2-GFP affect the behavior of the Arp2/3 complex in cells, we created diploid *S. pombe* strains with one copy of mutant Arp2-GFP and one copy of untagged wild-type Arp2 to keep the cells alive. While cells with untagged Y198F/T233A/T234A Arp2 built actin patches (Fig. 1 C; and Fig. 2 C), most of the GFP-tagged Arp2 with either the Y198F/T233A/T234A mutations (Fig. 1 E) or the lethal T233D/T234D mutations (Fig. 1 F) localized to the cytoplasm, with only a low level of fluorescence in patches. This explains why *S. pombe* strains with T233D/T234D Arp2 or Y198F/T233A/T234A Arp2-GFP mutations are not viable.

The lethal Arp2 mutants may render Arp2/3 complex nonfunctional by compromising its stability, rather than by modeling an altered phosphorylation state. Wild-type Arp2 dissociates from a fraction of fission yeast Arp2/3 complex during purification (Nolen and Pollard, 2008), and mutating the proposed phosphorylation sites at the interface between Arp2 and Arp3/ARPC4 could further weaken the binding of Arp2 to its neighbors. Unassembled components of Arp2/3 complex (Morrell et al., 1999) and other protein complexes, such as α- and β-spectrin (Woods and Lazarides, 1985) and WAVE regulatory complex (Kunda et al., 2003), are degraded. Consistent with degradation of unassembled Arp2, quantitative microscopy of diploid strains revealed lower cellular levels of mutant Arp2-GFP than wild-type Arp2-GFP (Fig. 1 G). Further, the Y198F/T233A/T234A Arp2/3 complex did not bind the GST-VCA column and elute intact from the subsequent ion exchange column during purification attempts (Fig. S1 F). The reduced growth of T233 mutants at 36°C is also consistent with compromised protein-protein interactions causing the observed defects, as increased temperature could weaken these interactions further. Therefore, our data suggest that mutations of Y198, T233 and T234 compromise some aspect of Arp2/3 complex function, reflected in the rate of actin patch assembly (Fig. 2).

### Effect of Y218 Arp2 mutations on actin patch assembly

Prior evidence indicates that the three proposed phosphorylation sites may not be the only residues at which phosphorylation regulates Arp2/3 complex (LeClaire et al., 2008). The *Drosophila* kinase NIK phosphorylated some sites on Arp2 and ARPC2 (detected by autoradiography with ^32^P) and restored the ability of phosphatase-treated *Acanthamoeba* Arp2/3 complex to nucleate actin filaments *in vitro* (LeClaire et al., 2015). However, NIK also partially restored the nucleation activity of Arp2/3 complex with Y202A/T237A/T238A Arp2, even though it could not phosphorylate these three residues of Arp2 (LeClaire et al., 2015). One way to reconcile these observations is that NIK activates Arp2/3 complex by phosphorylating an uninvestigated site on one of its subunits including Arp2 and ARPC2.

Mass spectrometry revealed a previously unidentified phosphorylation site on the fission yeast Arp2 subunit: the conserved residue Y218 (Fig. 3 A; and Fig. S1 D). Haploid fission yeast strains with Y218 replaced by either phenylalanine or glutamate were viable and built branched actin patches (Fig. 3). However, both blocking and mimicking phosphorylation of Y218 subtly decreased Arp2/3 complex activity (Fig. 3 H; and Fig. S2 H). Therefore, phosphorylation at Y218 is not essential for Arp2/3 complex to be active, and if the Y218E substitution effectively mimicked phosphotyrosine, then any effects due to mutations at this site resulted from compromising the protein structure. Future research should consider if phosphorylation at undetected sites on Arp2 and/or on another subunits activates Arp2/3 complex.

**Figure 3.**
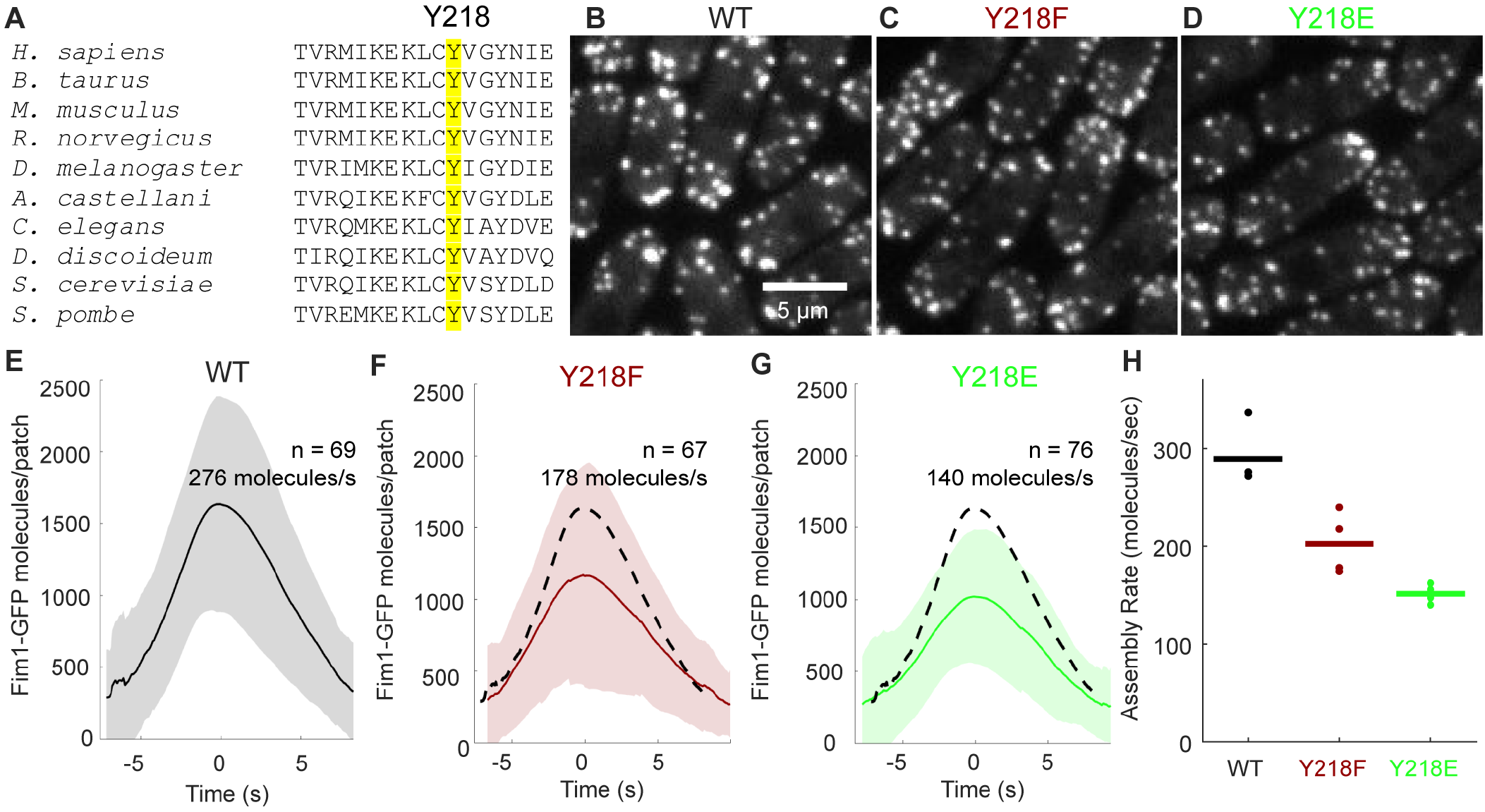
Phosphorylation at conserved Arp2 residue Y218 is not essential for Arp2/3 complex activity. **(A)** Sequence alignment of *S. pombe* Arp2 residues 208-224 with homologous regions of Arp2 in 9 other eukaryotes.Y218 and homologous residues are highlighted. **(B-D)** Sum projection confocal fluorescence images (6 Z-sections with 0.6 μm spacing) of haploid *S. pombe* endogenously expressing Fim1-GFP with **(B)** wild-type Arp2, **(C)** Arp2 Y218F or **(D)** Arp2 Y218E. **(E-G)** Mean numbers of Fim1-GFP molecules over time in 67-76 aligned actin patch tracks from **(E)** wild-type cells, **(F)** Y218F Arp2 or **(G)** Y218E Arp2. Dashed lines represent mean number of molecules per patch from wild-type Arp2 alignment in Panel E. **(H)** Rates of patch assembly for 3-4 replicates (24-76 patches per replicate) of wild-type and Y218 Arp2 mutants. Points indicate assembly rates of each replicate; horizontal bars denote the mean assembly rate.

## MATERIALS AND METHODS

### Purification of the Arp2/3 complex

Fission yeast was grown in 4-8 liters YE5S culture medium until the OD_595_ of a 1:5 dilution was 0.3-0.5 (1.5-2.5*10^7^ cells/ml), at which point a further 70 g/L YE5S powder was added and cells were allowed to resume growing. Cultures were pelleted and frozen when the OD_595_ of a 1:10 dilution was 0.55-0.65 (5.5-6.5*10^7^ cells/ml). Cells were thawed, lysed using a microfluidizer, centrifuged and Arp2/3 complex was purified from the soluble fraction by chromatography on a GST-VCA affinity column (GE Healthcare Life Sciences), a MonoQ 5/50 ion exchange column (GE Healthcare Life Sciences) and a HiLoad Superdex 200 16/60 gel purification column (GE Healthcare Life Sciences) (Ti et al., 2011). Roche cOmplete™ EDTA-free protease inhibitor tablets (Millipore Sigma) were added as the pellet thawed and periodically during initial centrifugation and dialysis steps. Roche PhosSTOP™ phosphatase inhibitor tablets (Millipore Sigma) were added alongside protease inhibitors in some preparations. The concentration of purified Arp2/3 complex was measured by absorption at 290 nm (ε = 139,030 M^−1^ cm^−1^).

### Mass spectrometry of *S. pombe* Arp2 subunit

Purity was confirmed by SDS-PAGE and staining the seven bands corresponding to the Arp2/3 complex subunits with Coomassie blue. The Arp2 (44 kDa) band was cut from the gel and digested with 6.67 μg/ml trypsin for 18 h at 37°C in 10 mM NH_4_HCO_3_ buffer. Digested samples were analyzed using liquid chromatography-tandem mass spectrometry (LC-MS/MS) on a Waters/Micromass AB QSTAR Elite mass spectrometer (Stone and Williams, 2009) by the Yale Mass Spectrometry and Proteomics Resource of the W.M. Keck Foundation. Phosphopeptides were identified using the Mascot algorithm (Hirosawa et al., 1993) through the Yale Protein Expression Database (Colangelo et al., 2015). Mass spectra from fragmentation of each identified phosphopeptide were checked (Fig. S1, C and D) to ensure that fragments identified were sufficient to distinguish phosphorylation from other chemical modifications.

### Generation of haploid and diploid *S. pombe* Arp2 mutant strains

We first employed a two-step homologous recombination process (Bahler et al., 1998; Grimm et al., 1988) to create point mutations in *arp2* of diploid *S. pombe* strains, because *arp2* is an essential gene. Haploid *S. pombe* strains with complementary adenine auxotrophy mutations *ade6-M210* and *ade6-M216* (Hoffman et al., 2016; Liang et al., 1999) were crossed to create diploid *S. pombe*, which we selected using adenine-deficient media. One copy of the wild-type *arp2* gene in diploid *S. pombe* lacking *ura4* was replaced by homologous recombination with a *ura4^+^* selectable marker, and transformed cells were selected on EMM5S-ura plates. The *ura4^+^* marker was replaced with mutant *arp2* or *arp2*-GFP adjacent to the *kan^R^* resistance marker, and transformants were selected on YE5S-G418 plates.

To create haploid yeast strains, we induced diploid yeast to sporulate on SPA5S medium for 24-48 h and then dissected tetrads on YE5S plates. Tetrads were grown for several days at 25°C and then replica plated onto YE5S-G418 plates to identify progeny with the *arp2* mutation and the *kan^R^* allele. All mutations were confirmed by DNA sequencing. A mutation was judged to be lethal if only two of four spores grew from each tetrad, corresponding to the two haploid progeny with the wild-type *arp2* allele. These progeny did not possess the *kan^R^* resistance allele and did not grow on YE5S-G418.

Haploid yeast strains were crossed with Fim1-GFP haploid yeast and tetrads dissected to create Fim1-GFP labeled haploid strains.

### Imaging *S. pombe* cells

*S. pombe* strains were grown in YE5S media in log phase (OD_595_<0.8) for 36 h. The cells were harvested at OD_595_ 0.4-0.7 (4-7*10^6^ cells/mL), washed twice with EMM5S media, and mounted on EMM5S 2% agarose pads. Images were acquired at room temperature with an Olympus IX-71 microscope with a 100x/NA 1.4 Plan Apo lens (Olympus) and an Andor CSU-X1 spinning disk confocal system with an iXON-EMCCD camera (Andor Technology).

Fluorescence was excited using a Coherent OBIS 488 nm LS 20 mW laser with adjustable power and an internal power meter. Time lapse movies of endocytic actin patches were acquired with Andor iQ2 software (Andor Technology); other images were acquired with the μManager 1.4 plugin (Stuurman et al., 2010) for ImageJ (Schneider et al., 2012).

### Generating the calibration curve and calculating number of molecules per cell

21 Z-sections with an 0.6 μm spacing were taken of eight yeast strains: one non-fluorescent strain (FY528) and seven endogenously expressing one of the following GFP-tagged proteins: Myo2p, Ain1p, Acp2p, ARPC5, Arp2p, Arp3p and Fim1p (Fig. S3 A; Wu and Pollard, 2005). Cells in sum projection images were automatically circled using the Mitotic Analysis and Recording System (MAARS) (Li et al., 2017) plugin for ImageJ using 21 bright-field Z-sections spaced 0.6 μm apart (Fig. S3 B). We corrected for autofluorescence by subtracting the total fluorescent intensity per cell in the non-fluorescent strain.

The number of fluorescent molecules per cell in each strain was previously obtained by quantitative immunoblotting (Wu and Pollard, 2005). We plotted this data against the total fluorescent intensity per cell in each strain (Fig. S3 C), which was calculated using an original ImageJ plugin. The slope of the best-fit line yielded the relationship between the total fluorescence intensity in an a given intracellular region and the number of fluorescent molecules it contained.

We employed the calibration curve to calculate the number of fluorescent molecules in each diploid strain. Autofluorescence was corrected by subtracting the total fluorescence intensity per cell in the diploid strain without any GFP (AE15D).

### Acquiring actin patch tracks

We took 3-4 time lapse movies with 1 s intervals for 60 s of each haploid yeast strain expressing endogenous Fim1-GFP. Each image consisted of 6 Z-sections separated by 0.6 μm collected with 5 mW laser power (as measured at the emission source; approximate power at the sample was 0.13 mW). An original MATLAB (MathWorks) script corrected the images for uneven illumination, camera noise and photobleaching (Fig. S3, D-I). Endocytic actin patches in corrected time lapse movies were identified and tracked using the Fiji (Schindelin et al., 2012) plugin PatchTrackingTools (Fig. S2, A and B; Berro and Pollard, 2014b). Identified patches were screened manually to reject records missing the beginning of assembly or the end of disassembly or where two patches overlapped.

All accepted patch tracks within each time lapse movie were aligned and their total fluorescence intensities (integrated density) were averaged using temporal super-resolution realignment (Fig. S2 C; Berro and Pollard, 2014a). The calibration curve was used to derive the number of molecules in each actin patch from its total fluorescence intensity. The resulting number of Fim1-GFP molecules in the average actin patch was plotted as a function of time, generating one assembly/disassembly curve per time lapse movie. The movie with the largest sample of patches was chosen for example figures.

We used an original MATLAB script to fit straight lines to the most linear 3.7 s regions of the assembly and disassembly phases to measure these rates (Fig. S2, C and D). The mean assembly rate for each strain was determined by averaging the assembly rates observed for 3-4 averaged assembly/disassembly curves (Fig. S2 E). Strains were plotted in MATLAB using colorblind-safe colors (Martin Krzwinski, http://mkweb.bcgsc.ca/biovis2012/color-blindness-palette.png). Shaded regions indicating SD were generated using the shadedErrorBar MATLAB script (Rob Campbell, https://www.mathworks.com/matlabcentral/fileexchange/26311-raacampbell-shadederrorbar).

### Automatic photobleaching correction of yeast time lapse movies

We automated the photobleaching correction process with an original MATLAB function. Time lapse movies were corrected for camera noise and uneven illumination. Poorly illuminated areas were identified and cropped using the best-fitting two-dimensional Gaussian function to the uneven illumination correction image. To identify intracellular regions, each of the first ten frames of each time lapse movie was thresholded by brightness. Pixels brighter than the threshold in eight out of ten frames were assumed to be intracellular (Fig. S3 F).

To determine the threshold used, all pixels in the cropped image were binned according to brightness. The resulting histogram consisted of two populations: one of intracellular and one of background pixels. Both populations were expected to have a normal brightness distribution, and the histogram was therefore fit to a two-term Gaussian function. Background pixels were identified by fitting a one-term Gaussian function to the region of the histogram preceding the peak. The threshold was set at the intensity bin for which 95% of pixels were predicted to be intracellular (Fig. S3 E).

The median fluorescence of the intracellular pixels was calculated for each frame of the time lapse movie, plotted and fit to a double exponential decay function, where the first exponential term corresponds to bleaching of autofluorescence and the second likely represents photobleaching of GFP (Fig. S3 G). The decay function was normalized to start at 1, and time lapse movies were corrected by dividing each frame by the corresponding value of the normalized decay function (Fig. S3 I).

### *S. pombe* growth assays

Cells were grown in YE5S for 36 h in log phase. Cultures were harvested at an OD_595_ of 0.4-0.6 (35-65×10^5^ cells/mL) and diluted to OD_595_ 0.1. After 1:10 serial dilutions, 10 μL samples were plated on YE5S and incubated for 24-48 h at 32°C or 36°C (Fig. S1 E).

### Online supplemental material

Fig. S1 (A and B) shows the location of the previously proposed phosphorylation sites Y198, T233 and T234 on Arp2 and that they are widely conserved in eukaryotes. Fig. S1 (C and D) depicts mass spectra from fragmentation of peptides containing phosphorylation at Y198 and Y218. Fig. S1 (E-G) lists generated Arp2 mutations at Y198, T233, T234 and Y218 and shows the effect of some mutations on *S. pombe* growth and on purification of the Arp2/3 complex by binding to a GST-VCA affinity column. Fig. S2 (A-E) shows tracking and temporal super-resolution realignment of Fim1-GFP labeled actin patches, (F) lists Fim1-GFP *S. pombe* strains, and (G and H) contains the assembly rate data obtained through the patch tracking program. Fig. S3 (A) lists the strains employed for the calibration curve, and (B and C) shows how images are segmented and the total intracellular fluorescence of each strain is used to give the calibration curve. Fig. S3 (D-I) demonstrates photobleaching correction of a sample actin patch movie.

## Acknowledgments

Research reported in this publication was supported by National Institute of General Medical Sciences of the National Institutes of Health under award number R01GM026338. The content is solely the responsibility of the authors and does not necessarily represent the official views of the National Institutes of Health. This research was also supported by fellowships to Alexander Epstein from the Arnold and Mabel Beckman Foundation and from the Yale University Office of Science and Quantitative Reasoning. The authors thank Dr. Julien Berro for help with the with automated patch tracking and analysis software, Dr. Tong Li for support with the MAARS segmentation software, Helen Sun for reading the manuscript, Jean Kanyo for technical assistance with mass spectrometry, and Dr. Samantha E.R. Dundon for invaluable assistance with yeast culture techniques, microscopy and editing.

The authors declare no competing financial interests.

## Author contributions

A.E.E, S.E.-S. and T.D.P. designed research; A.E.E and S.E.-S performed the research; A.E.E, S.E.-S. and T.D.P. analyzed the data; A.E.E and T.D.P. wrote the paper.

## Abbreviation List

NPF: nucleation promoting factor
WASp: Wiskott-Aldrich syndrome protein
MD simulations: molecular dynamics simulations
NIK: Nck-interacting kinase
LC-MS/MS: liquid chromatography-tandem mass spectrometry

## Supplemental materials

**Figure S1.**
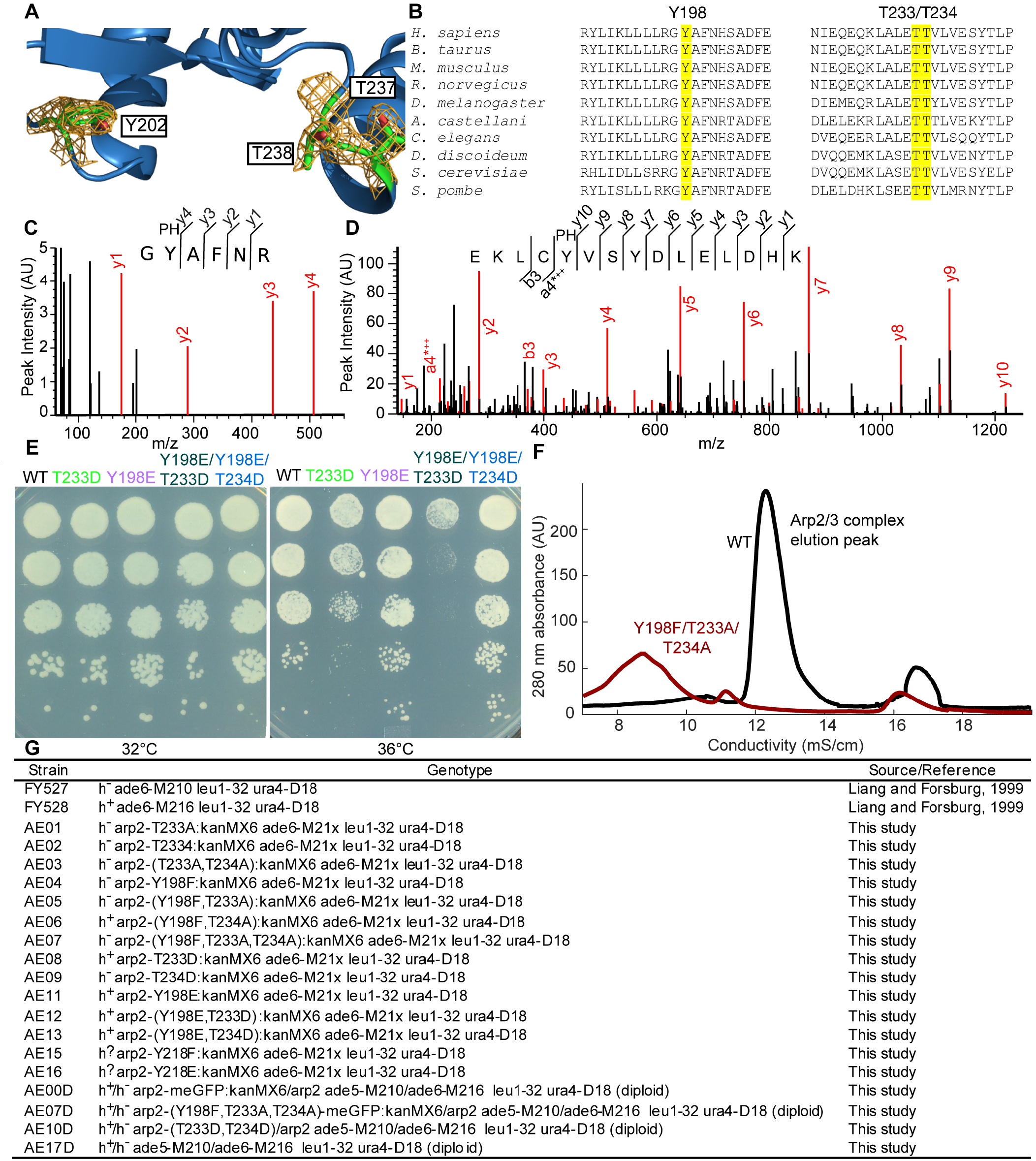
Proposed phosphorylation sites on Arp2. **(A)** Ribbon diagram of *Bos taurus* Arp2 backbone (blue) with stick models (green) of Y202, T237 and T238. The orange mesh is the electron density map (contour level 2.5 σ), which reveals no density corresponding to a phosphate group at any of the oxygens (red) of the three proposed phosphorylation sites (Robinson et al., 2001; pdb: 1k8k). **(B)** Sequence alignment of *S. pombe* Arp2 residues 187-207 and 222-245 with homologous sequences in 9 other eukaryotes. Y198, T233 and T234 and homologous residues are highlighted. **(C and D)** Mass spectra from fragmentation of **(C)** Y198 and **(D)** Y218 phosphopeptides identified using LC-MS/MS of Arp2 from Arp2/3 complex purified in the presence of phosphatase inhibitors. Red peaks were identified by the Mascot algorithm as associated with a fragment of the phosphopeptide; selected fragments are labeled on both the spectrum and the peptide sequence. y(*n*)-peaks represent *n*-residue C-terminal fragments; a/b(*n*) peaks represent *n*-residue N-terminal fragments. * indicates loss of water and + represents positive charge. **(E)** Growth of 1:10 serial dilutions of wild-type cells and strains with phosphomimetic Arp2 mutations on YE5S plates at 32°C and 36°C. **(F)** Analysis of elution fractions from anion exchange chromatography of Arp2/3 complex after elution from the GST-VCA affinity column. No protein from the Y198F/T233A/T234A strain elutes at the position of wild-type Arp2/3 complex. **(G)** Table of viable haploid and diploid strains with mutations at Y198, T233, T234 and Y218.

**Figure S2.**
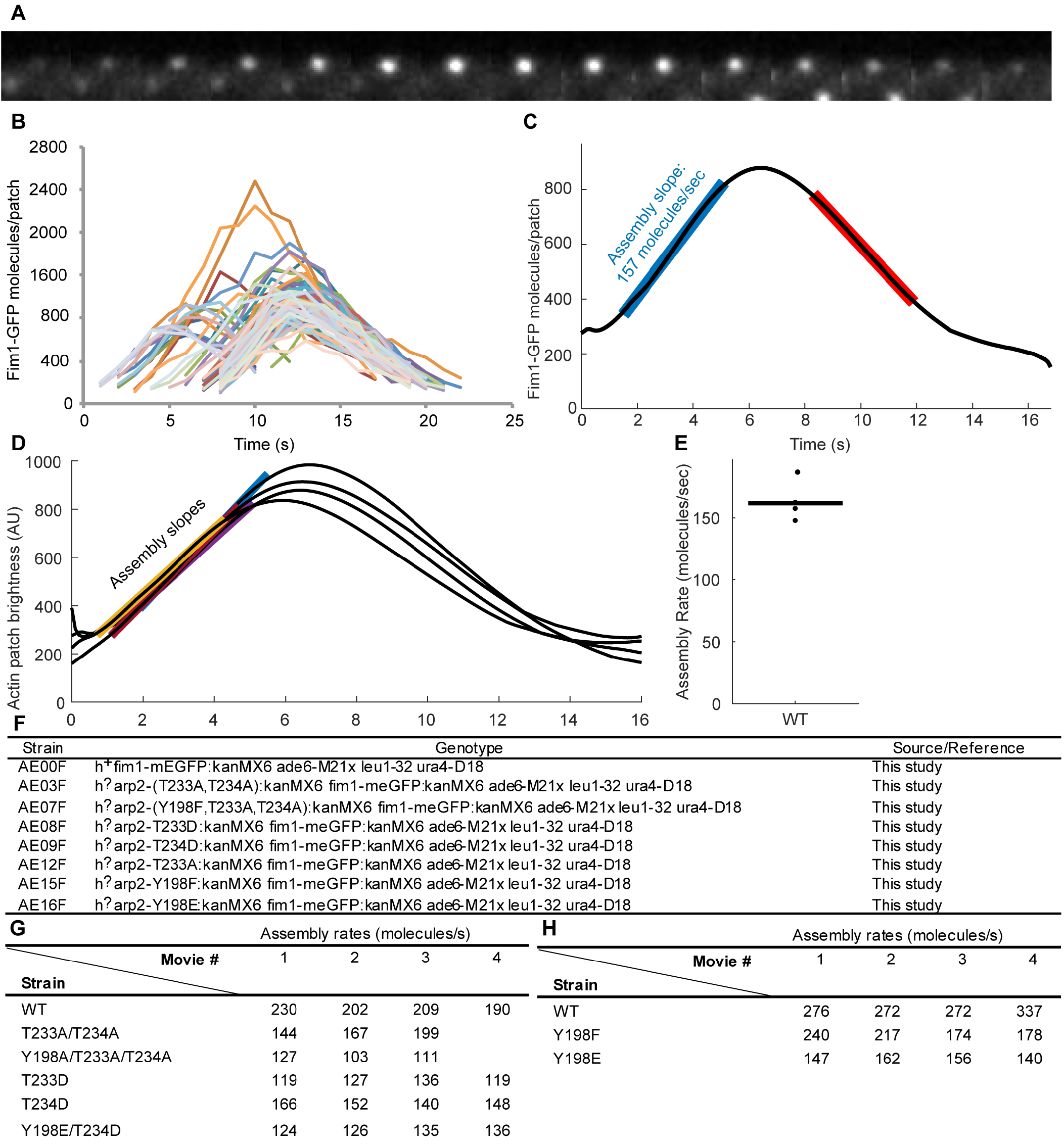
Time course of Fim1-GFP appearance and disappearance in actin patches. **(A)** Time series of fluorescence micrographs at 1 s intervals of an individual actin patch in a cell expressing Fim1-GFP reconstructed from sum projections of 6 Z-sections. **(B)** Numbers of Fim1-GFP molecules over time in 95 individual actin patches in wild-type cells. **(C)** Mean number of Fim1-GFP molecules per actin patch over time after continuous realignment of patches in panel B. The assembly phase (blue) and disassembly phase (red) are highlighted. **(D)** Aligned Fim1-GFP actin patch tracks from four separate time lapse movies of wild-type *S. pombe*, with assembly phases highlighted. **(E)** Assembly rates from the four time lapse movies in Panel D, with bar representing the mean. **(F)** Table of Fim1-GFP *S. pombe* strains employed for patch tracking analysis. **(G and H)** Assembly rates observed for 3-4 movies of wild-type and mutant Fim1-GFP *S. pombe* strains.

**Figure S3.**
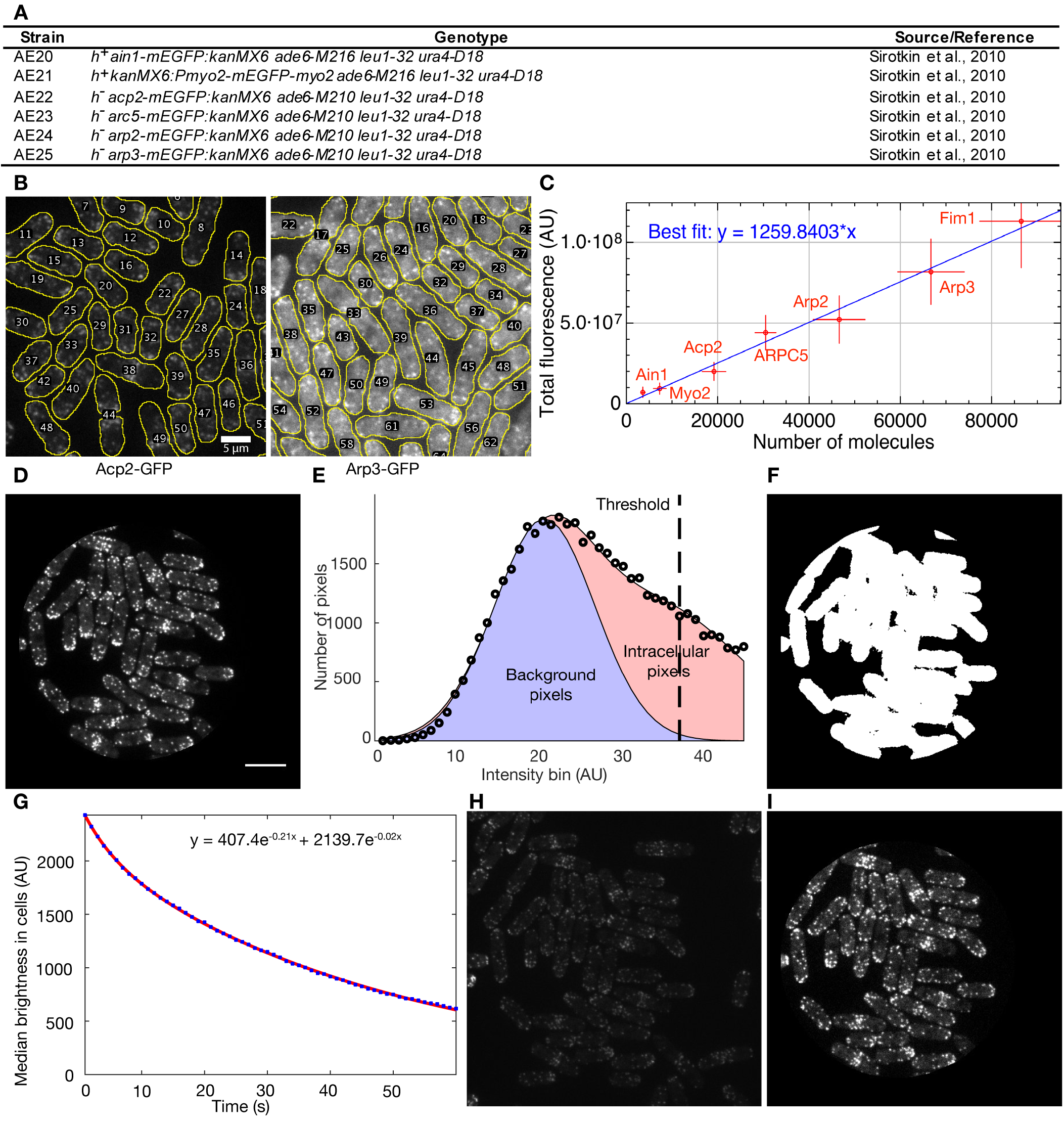
Calibration curve and automated photobleaching correction for analysis of actin patches in time-lapse movies. **(A)** Table of *S. pombe* strains used for the calibration curve. **(B)** Example segmentations using MAARS software of fluorescence micrographs of fission yeast cells expressing Acp2-GFP or Arp3-GFP. These were two of the seven *S. pombe* strains used to construct the calibration curve. **(C)** Calibration curve constructed using seven endogenously expressed GFP-tagged proteins. Numbers of molecules per cell were published (Wu and Pollard, 2005). **(D)** First frame of a sample time lapse movie consisting of six optical sections with 0.6 μm spacing, cropped to remove areas of highly uneven illumination. **(E)** Determination of segmentation threshold using Gaussian fits to pixel brightness histogram. Circles: histogram of pixel brightness within uncropped area of first frame. Blue area: Gaussian fit to section of pixel brightness histogram preceding the peak. Red area: Two-term Gaussian fit to entire pixel brightness histogram. Dashed line: Brightness threshold used for segmentation, at which 95% of pixels are thought to be intracellular. **(F)** Binary segmentation of first 10 frames of the sample time lapse movie to highlight intracellular regions. **(G)** Median pixel brightness within intracellular region over time, fit with a double-exponential function. **(H and I)** Last frame of the sample time lapse movie **(H)** before and **(I)** after cropping and photobleaching correction.

